# Long-term Locus Coeruleus Stimulation Exacerbates Tau Pathology in PS19 Mice

**DOI:** 10.1101/2025.11.01.686018

**Authors:** Yuhan Nong, Steven Wellman, Hong Zhang, Yuxiang (Andy) Liu, Elentina K. Argyrousi, Ottavio Arancio, Qi Wang

## Abstract

**Background:** Alzheimer’s disease (AD) is the most common form of dementia, characterized by the accumulation of amyloid-β (Aβ) plaques and hyperphosphorylated Tau tangles. The locus coeruleus (LC) is among the first brain regions to show degeneration and Tau pathology during the early stages of AD. Previous studies have demonstrated that short-term chemogenetic LC stimulation can improve memory performance in the TgF344-AD rat model, while long-term norepinephrine (NE) reuptake inhibition can worsen memory deficits in the ADLP^Tau^ mouse model. However, the effects of long-term LC stimulation in Tau mouse models on memory, synaptic plasticity, and tauopathy remain unclear.

**Objective:** To evaluate the impact of long-term locus coeruleus stimulation on memory, synaptic plasticity, and tauopathy in PS19 mice using behavioral paradigms, electrophysiological recordings, and immunohistochemical analysis.

**Methods:** The radial arm water maze and fear conditioning test were conducted to assess memory performance in PS19 mice with and without long-term LC stimulation. Hippocampal long-term potentiation was recorded to evaluate the effect of long-term LC stimulation on synaptic plasticity. Immunohistochemistry was employed to examine Tau phosphorylation, neurodegeneration, and neuroinflammation.

**Results:** Long-term LC stimulation in PS19 mice exacerbated spatial memory deficits in the water maze, impaired contextual fear memory, reduced hippocampal LTP, and increased AEP expression, Tau hyperphosphorylation, and astrocyte activation.

**Conclusion:** Long-term LC stimulation may exacerbate memory deficits in PS19 mice by impairing synaptic plasticity and increasing neural degeneration in the hippocampus. Elevated norepinephrine levels resulting from long-term LC stimulation may increase AEP expression, contributing to Tau hyperphosphorylation in the LC.

## Introduction

Alzheimer’s disease (AD) is the leading cause of dementia, which is defined by the accumulation of amyloid beta (Aβ) and tau pathology in the brain (Self and Holtzman, 2023). Although substantial efforts have been made to develop treatments for AD, no effective therapy has emerged, possibly because most research has focused on the late stages of the disease, when significant memory impairment and physiological pathology have already occurred. This has led to a growing interest in investigating the early stages of AD, with the aim of uncovering strategies that could prevent or slow the initial development of the disease. In this work, we focus on the effect of long-term stimulation of the locus coeruleus (LC) on tau pathology as the LC is among the earliest brain regions to show hyperphosphorylated tau, long before the onset of memory deficits (Braak et al., 2006, 2011; Weinshenker, 2018; Mercan and Heneka, 2022).

The LC, located in the pons of the brainstem, is the brain’s principal source of norepinephrine (NE) (Poe et al., 2020; Slater and Wang, 2021). It sends widespread projections throughout the central nervous system and plays a crucial role in regulating arousal, attention, and the stress response (Benarroch, 2009; Kelberman et al., 2024). Given its broad influence on neural activity through different subtypes of adrenergic receptors, maintaining proper NE signaling from the LC is essential for normal cognitive and behavioral function (Rodenkirch et al., 2019; Jordan and Keller, 2023; Stanley et al., 2023; Ghosh and Maunsell, 2024; Grimm et al., 2024; Osorio-Forero et al., 2024; Nong et al., 2025). In AD, the LC undergoes significant degeneration, as observed in both postmortem human tissue and AD mouse models (Braak et al., 2006, 2011). Preserving LC integrity has thus become a focus of therapeutic interest. Various drugs have been developed to support NE levels in the brain (Gutiérrez et al., 2022). In addition, research has shown that short-term chemogenetic activation of the LC can improve memory performance in a rat model expressing both presenilin-1 (PS1ΔE9) and mutant amyloid precursor protein (APPsw) (Rorabaugh et al., 2017). However, emerging evidence suggests that excessive NE may have harmful effects in the context of AD. Chronic stress, which elevates brain NE levels, can exacerbate Aβ accumulation, tau hyperphosphorylation, and cognitive deficits in AD models (Finlay et al., 1995; Sotiropoulos et al., 2011; Lyons and Bartolomucci, 2020). Furthermore, elevated NE levels have been reported in some AD patients (Gannon and Wang, 2019). A recent study demonstrated that prolonged administration of the NE reuptake inhibitor reboxetine (RBX) induced memory impairment and neurodegeneration in tauopathy mouse models (Jeong et al., 2025). This may be due to the NE metabolite 3,4-dihydroxyphenyl glycolaldehyde (DOPEGAL), which activates asparagine endopeptidase (AEP), leading to tau hyperphosphorylation and neuronal loss (Zhang et al., 2014). Additionally, DOPEGAL has later been shown to bind directly to tau and promote its aggregation (Kang et al., 2020, 2022).

To explore the therapeutic potential of treatments targeting the NE system in AD, a deeper understanding of NE’s role in AD pathogenesis is essential. Given that the LC is the first region to exhibit tau pathology, we hypothesized that early and sustained activation of LC neurons may influence disease progression. Therefore, we initiated long-term LC stimulation in PS19 mice starting at 3 months of age, before the appearance of pathological features, and continued the intervention in the following 3 months. To achieve this, we employed a chemogenetic approach using excitatory DREADDs targeting dopamine β-hydroxylase (DbH)-expressing neurons in the LC. Mice received daily administration of DCZ/CNO on weekdays over a three-month period. Following the chronic LC stimulation, we assessed the animals’ cognitive function using the radial arm water maze and fear conditioning tests. Subsequently, we performed in vitro long-term potentiation (LTP) recordings to assess the effects of chronic LC stimulation on synaptic plasticity. Furthermore, we utilized immunohistochemical analyses to evaluate key pathological features of tauopathy, including tau accumulation, neurodegeneration, and neuroinflammation.

## Materials and Methods

### Animals

All procedures were approved by the Columbia University Institutional Animal Care and Use Committee (IACUC) and conducted in accordance with NIH guidelines. Male DbH-Cre mice (The Jackson Laboratory, Cat #: 033951) were crossed with female PS19 mice (The Jackson Laboratory, Cat #: 008169) to generate double transgenic PS19xDbH-Cre offspring. 50 mice (27 females), consisting of double-positive PS19xDbH-Cre mice (n = 19, 10 females), single-positive PS19 mice (n=17, 9 females), and wild-type littermates (n=14, 8 females), were used in the study (**Fig. 1a**). Mice were housed under a 12-hour light/dark cycle with ad libitum access to food.

**Figure 1.**
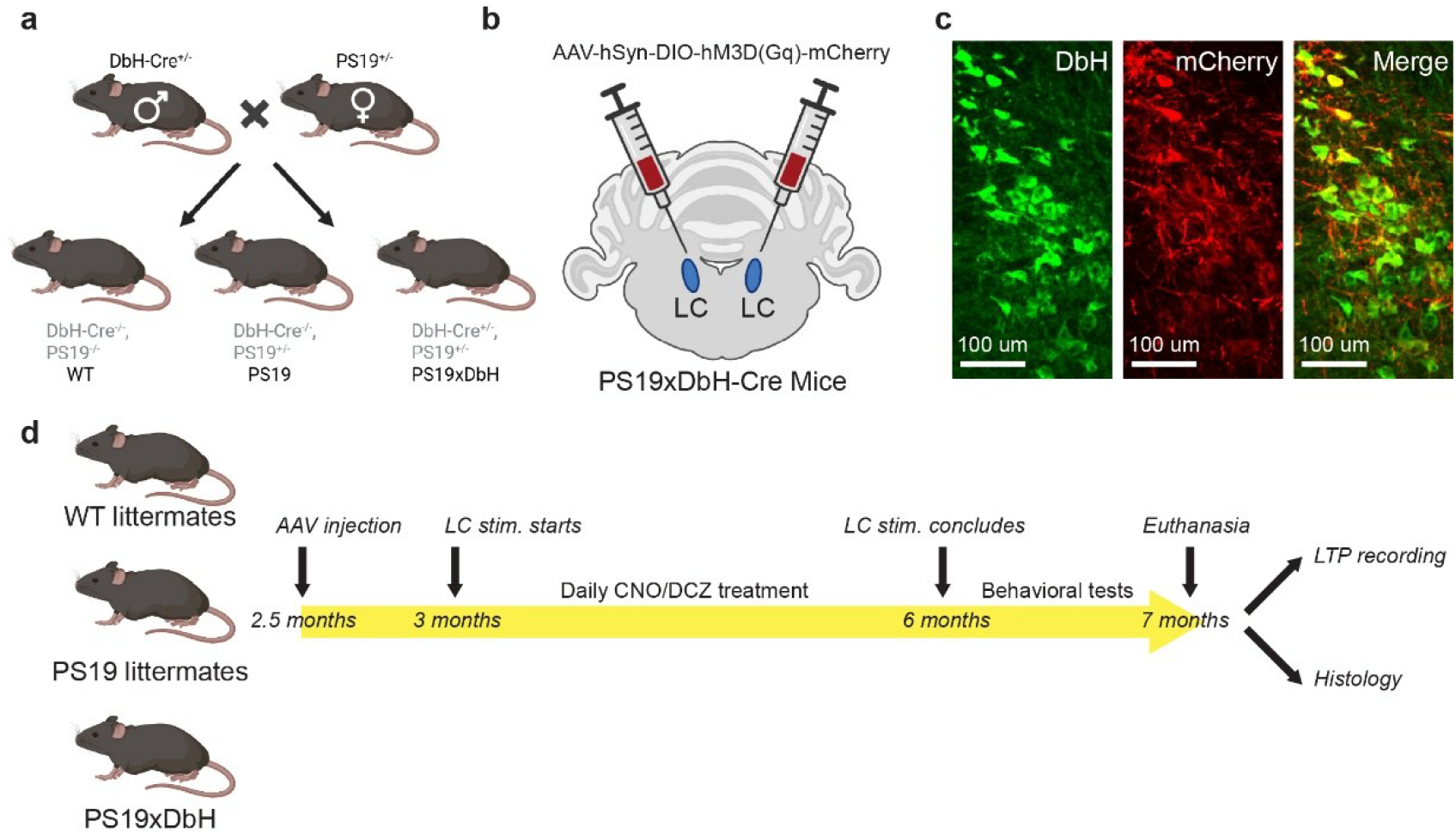
Experimental setup. **a)** Breeding paradigms to generate experimental animals for the study. **b)** Diagram of AAV-mediated expression of hM3D(Gq) DREADD receptors in the LC. **c)** Histological confirmation of selective expressions of the DREADD receptors in LC neurons. **d)** Timeline of experiments.

### Stereotaxic surgery

The surgical procedure for injecting adeno-associated viral (AAV) vector is the same as described previously (Weiss et al., 2024, 2025; Apte et al., 2025). Briefly, mice were anesthetized with isoflurane (5% for induction, 1-2% for maintenance) and secured in a stereotaxic frame (Kopf Instrument, CA). Body temperature was maintained at 37 °C using a feedback-controlled heating pad (FHC, Bowdoinham, ME). Once the animal reached a surgical anesthesia stage, lidocaine hydrochloride and buprenorphine (0.05 mg/kg) were administered subcutaneously prior to making the incision. After removing the fur over the scalp, the incision site was cleaned with alternating scrubs of betadine and 70% alcohol. To inject AAV vector into the LC, small burr holes were drilled bilaterally above the LC (AP: −5.3 mm, ML: ±0.9 mm relative to bregma), and sterile saline was applied to prevent cortical drying. Pulled glass capillary micropipettes (Drummond Scientific, Broomall, PA) were back-filled with AAV solution (pAAV-hSyn-DIO-hM3D(Gq)-mCherry, Addgene #: 44361) and connected to a precision injection system (Nanoliter 2020, World Precision Instruments, Sarasota, FL). The AAV solution was injected at three DV depths each side (−2.9 mm, −3.0 mm, −3.1 mm, respectively. ∼200 nL per site, 1 nL/s) to ensure full LC coverage. After each injection, the pipette was left in place for at least 10 minutes to minimize backflow before being slowly withdrawn. The burr holes were sealed with sterile bone wax, and the wound were closed with sutures. Baytril (5 mg/kg, S.C.) and Ketoprofen (5 mg/kg, S.C.) were given immediately after the procedure and subsequently four additional times over the following days. Animal body weight was monitored daily for five consecutive days.

### Drug Administration

To achieve long-term stimulation of LC neurons, we employed a chemogenetic approach by first selectively expressing DREADDs (Designer Receptors Exclusively Activated by Designer Drugs) in LC neurons in a Cre-dependent manner (**Fig. 1b&c**). DREADD agonists clozapine-N-oxide (CNO, 2 mice) or deschloroclozapine (DCZ, 13 mice; no significant difference was found between CNO- and DCZ-treated mice) was subsequently administered daily on weekdays to selectively activate LC neurons for 3 months (**Fig. 1d**). To avoid potential stress and injury, such as esophageal perforation and tracheal injury, associated with long-term oral gavage, CNO (5 mg/kg) or DCZ (0.6 mg/kg) were dissolved in 5% sucrose solution and administered through drinking water (Oyama et al., 2022). To facilitate the administration of CNO/DCZ, all mice were mildly water-restricted and given a 30-minute *ad lib* access to water each day, during which drug-containing sucrose solution was first provided and mice were closely monitored to ensure complete consumption of the drug solution. In our experience, the sucrose solution containing CNO or DCZ was typically consumed within seconds. Ad lib water access was then provided for 30 minutes. CNO/DCZ administration was stopped for at least for 5 days before behavioral assays to avoid their possible short-term effects on the tasks.

### Behavioral studies

#### Radial arm water maze test

The radial arm water maze (RAWM) was conducted in a 120 cm diameter white circular pool filled with opaque water, achieved by adding non-toxic white paint (Acquarone et al., 2019). The water temperature was consistently maintained at 24 ± 2 °C throughout testing. A six-arm radial insert was placed in the center of the pool, forming six equally spaced arms extending outward. Visual cues were affixed to the walls surrounding the maze to assist spatial navigation. A circular escape platform (10 cm in diameter) was submerged at the end of one fixed goal arm. The platform location remained the same for each mouse across all trials, while the entry arm varied randomly. Mice were tested over two consecutive days with 15 trials each day. On day one, mice underwent a training protocol consisting of 15 trials, where the first 12 trials alternated between visible (flagged) and hidden (submerged 1 cm below the water surface) platforms. The final 3 trials on day one and all 15 trials on day two were conducted with the platform hidden.

Each trial lasted up to 60 seconds, during which the mouse was allowed to freely search for the platform. Once it found the platform, the mouse remained there for 15 seconds to observe the surrounding cues. If the mouse did not find the platform within 60 seconds, the experimenter guided it to the platform for the same 15-second observation period. Errors were recorded when the mouse entered a non-goal arm (defined as all four paws inside an arm) or failed to make a directional choice within 10 seconds. After each trial, the mouse was removed, gently dried, and returned to its home cage, placed under a heat lamp. To minimize fatigue and avoid overtraining, mice were tested in cohorts of 4–5 animals. Trials were interleaved across cohorts to space out testing throughout the day. Behavioral performance was analyzed in blocks of 3 consecutive trials, yielding a total of 10 blocks across the 30 trials.

#### Visible platform test

The visible platform test was used to evaluate potential visual, motor, or motivational impairments. It was conducted in the same circular pool used for the RAWM task. The test was carried out over two consecutive days, with mice undergoing two sets of trials per day. Each set consisted of three trials in which the mouse was trained to locate a visible escape platform, marked by a bottle cap placed on top. During each set, the platform was positioned in one of three quadrants of the pool, and the mouse was released from a fixed starting point corresponding to that platform location. Mice were gently placed into the water facing the wall, and each trial continued until the platform was found or a maximum of 60 seconds had elapsed. After each trial, if the mouse had not located the platform, it was guided to the platform and allowed to remain there for 15 seconds to observe spatial cues. Latency to find the platform and swimming speed were recorded and analyzed using a video tracking system (EthoVision XT, Noldus). Results were grouped into four blocks, with each block representing the average performance across one set of three trials.

#### Fear conditioning test

The fear conditioning test was used to assess associative fear memory in mice and was conducted over three consecutive days. On day one, mice were placed into a fear conditioning chamber (Noldus) and allowed to explore for 2 minutes before hearing a tone (2880 Hz, 85 dB), which served as the conditioned stimulus. During the final 2 seconds of the tone, a foot shock (0.8 mA) was delivered as the unconditioned stimulus. After the pairing, mice remained in the chamber for an additional 30 seconds without any stimuli. On day two, a contextual conditioning test was performed, in which the mice were returned to the same chamber for 5 minutes with no tone or shock. Freezing behavior, defined as the absence of all movement except for respiration, was recorded using an automated tracking system (EthoVision XT, Noldus). Mice showing freezing levels higher than 90% during the contextual test were excluded from analysis to avoid possible effects of motor deficits in PS19 mice or anxiety -like behaviors unrelated to associative learning. On day three, a cued conditioning test was performed, in which the mice were placed in a novel context created by altering the chamber’s walls, floor, and introducing a vanilla scent. The session lasted 5 minutes, with the first 2 minutes for free exploration followed by a 3-minute presentation of the tone. Freezing behavior during this period was used to assess cue-associated memory.

#### Sensory threshold assessment

The sensory threshold test was used to evaluate the mice’s perception of foot shocks to ensure that the results of the fear conditioning tests were not confounded by impaired perception of foot shocks. This test was performed on the final day of behavioral experiments, using the same chamber as the fear conditioning test. Mice were exposed to a series of 1-second foot shocks with increasing intensity, starting at 0.1 mA and rising by 0.1 mA every 30 seconds until reaching 0.7 mA. Behavior was recorded using a video tracking system (EthoVision XT, Noldus) and manually scored. The average shock intensity required to elicit the first visible response (flinching), the first major motor response (jumping), and the first audible response (vocalization) was calculated and presented in the results.

#### Open field test

The open field test was used to assess exploratory behavior and anxiety-like responses. Mice were placed in a novel square arena (30 × 30 × 50 cm) constructed from glass and visually isolated using white opaque coverings on all sides. Each mouse was allowed to freely explore the arena for 10 minutes, and the test was conducted over two consecutive days. Behavior was automatically recorded and analyzed using a video tracking system (EthoVision XT). The total distance traveled, percentage of time spent in the center zone, and number of center zone entries were quantified.

### Electrophysiological recordings

Mice were sacrificed by cervical dislocation, and the hippocampus was rapidly extracted following decapitation. Transverse hippocampal slices (400 μm thick) were prepared using a tissue chopper and immediately transferred to a recording chamber. Slices were continuously perfused with artificial cerebrospinal fluid (ACSF) bubbled with a gas mixture of 95% O_2_ and 5% CO_2_ to maintain physiological conditions. The ACSF contained (in mM): 124.0 NaCl, 4.4 KCl, 1.0 Na_2_HPO_4_, 25.0 NaHCO_3_, 2.0 CaCl_2_, 2.0 MgCl_2_, and 10.0 glucose. Slices were allowed to recover for at least 90 minutes before recordings. A bipolar tungsten stimulating electrode was positioned in the Schaffer collateral fibers, and a glass recording electrode filled with ACSF was placed in the stratum radiatum of the CA1 region. Input-output curves were generated to determine the maximum evoked slope, and baseline synaptic transmission was recorded every minute at ∼35% of the maximal slope. Once a stable baseline was established for 30 minutes, long-term potentiation (LTP) was induced using a theta-burst stimulation protocol: four pulses at 100 Hz repeated at 5 Hz, with each tetanus consisting of three 10-burst trains delivered 15 seconds apart. Field excitatory postsynaptic potentials (fEPSPs) were recorded for 2 hours following stimulation. LTP magnitude was quantified as the fEPSP slope, normalized to the baseline, and results were expressed as mean ± SEM.

#### Immunohistochemistry

For immunohistochemistry, only mice that did not undergo LTP recordings were used. At the conclusion of the study, these mice were transcardially perfused with PBS, followed immediately by ice-cold 4% paraformaldehyde (PFA). Brains were carefully extracted and post-fixed in 4% PFA at 4 °C overnight, then cryoprotected in 30% sucrose (w/v in PBS) at 4 °C for three days. After cryoprotection, brains were embedded in Optimum Cutting Temperature (OCT) compound, and 25-μm coronal sections containing the LC were obtained using a cryostat (Leica CM 1950). Brain slices were then washed three times in PBS and incubated for 2 hours at room temperature in a blocking solution containing 10% normal donkey serum and 1% Triton X-100 in PBS (Liu et al., 2025). Following blocking, slices were washed six times, alternating between PBS with 0.1% Tween-20 and PBS alone. Slices were then incubated for 48 hours at 4°C with the primary antibody solution: 1:500 rat anti-mCherry (M11217, Invitrogen), 1:1000 chicken anti-tyrosine hydroxylase (TYH-0020, Aveslabs) for TH detection, 1:500 rabbit anti-asparagine endopeptidase (93627S, Cell Signaling Technology) for AEP detection, 1:500 rabbit anti-dopamine ß hydroxylase (AB209487, Abcam) for DbH detection, 1:250 biotin anti-AT8 (MN1020B, Invitrogen) for pTau detection, 1:500 mouse anti-NeuN (MA5-33103, Invitrogen) as a neuronal marker and 1:500 chicken anti-GFAP (AB4674, Abcam) for GFAP detection.

Following primary antibody staining, the slices were washed three times in PBS, followed by 5 hours of incubation with the appropriate secondary antibodies. We used 1:500 Alexa Fluor 568-conjugated goat anti-rat (A11077, Invitrogen) to amplify mCherry. For staining DbH, AEP, we used 1:500 Alexa Fluor 488-conjugated Donkey anti-rabbit (AB150061, Abcam). For staining TH and GFAP, we used 1:500 Alexa Fluor 647-conjugated Donkey anti-chicken (703-605-155, Jackson Immuno Research). For staining AT8, we used 1:500 Alexa Fluor 647-conjugated Streptavidin (016-600-084, Jackson Immuno Research). For staining NeuN, we used 1:500 Alexa Fluor 568-conjugated Donkey anti-mouse (AB175700, Abcam). Afterward, sections were washed three times in PBS, then mounted using Fluoromount-G with DAPI (00-4959-52, ThermoFisher).

#### Image processing and quantification

To evaluate the level of Tau pathological effects, neurodegeneration and inflammatory response in the brain regions of our interest, each slice was imaged using Z-stack and tile scanning with a confocal microscope (Nikon Ti2) equipped with a Yokogawa CSU-W1 spinning disk. Regions of interest (ROIs) encompassing the full extent of hippocampus (AP ∼-2.0 mm) or LC (AP ∼-5.3 mm) were manually drawn on low-magnification (10x) pre-scanned images based on DAPI, Dbh or TH signals and referencing the mouse brain atlas. For each slice, a composite image was generated by projecting all stacks using their maximum intensity. One slice per animal was analyzed for each of AT8, AEP, GFAP and NeuN staining. Images were blinded, a threshold was determined across all images for each type of staining, and contours of immunoreactivity were selected based on the threshold (Chalermpalanupap et al., 2018). Area of contour was then calculated using the “Measure” feature of ImageJ, which was used to further calculate the percentage of co-expression (area of immunoreactivity within nuclei of interest / total area of nuclei *100%). For quantifying the expression of AT8 or AEP among LC neurons, we calculated the area of immunoreactivity normalized by the total Dbh+ or TH+ area. For quantifying the expression of NeuN or GFAP among hippocampus neurons, we calculated the area of immunoreactivity normalized by the total DAPI+ area.

## Results

### Chronic LC stimulation exacerbated the memory performance of PS19 mice

To assess spatial working memory, we employed a two-day radial arm water maze (RAWM) test, which requires short-term reference memory (Acquarone et al., 2019). Compared to wild-type (WT) littermates and PS19 mice, PS19xDbH mice with 3-month LC-stimulation failed to learn the task and exhibited significantly more errors (p < 0.05) (**Figure 2a**). Control experiments using the visible platform test excluded potential confounding factors such as deficits in visual, motor, or motivational abilities (Puzzo et al., 2009), as all groups displayed comparable swimming speeds (p=0.16, one way ANOVA test. **Fig. 2b**) and latencies to locate the platform (p =0.66 for session 1, p=0.58 for session 2, p=0.78 for session 3, and p=0.28 for session 4, one way ANOVA tests. **Fig. 2c**). Additionally, to rule out potential direct effects of DCZ/CNO as compared to their effects on DREADD receptors, we administered sucrose water (i.e. vehicle for DCZ/CNO administration) to AAV-injected PS19xDbH mice. The performance of this cohort of mice did not differ from that of saline-injected PS19 littermate controls, suggesting that the impaired performance was primarily due to chronic activation of the LC (**Fig. 2a**).

**Figure 2.**
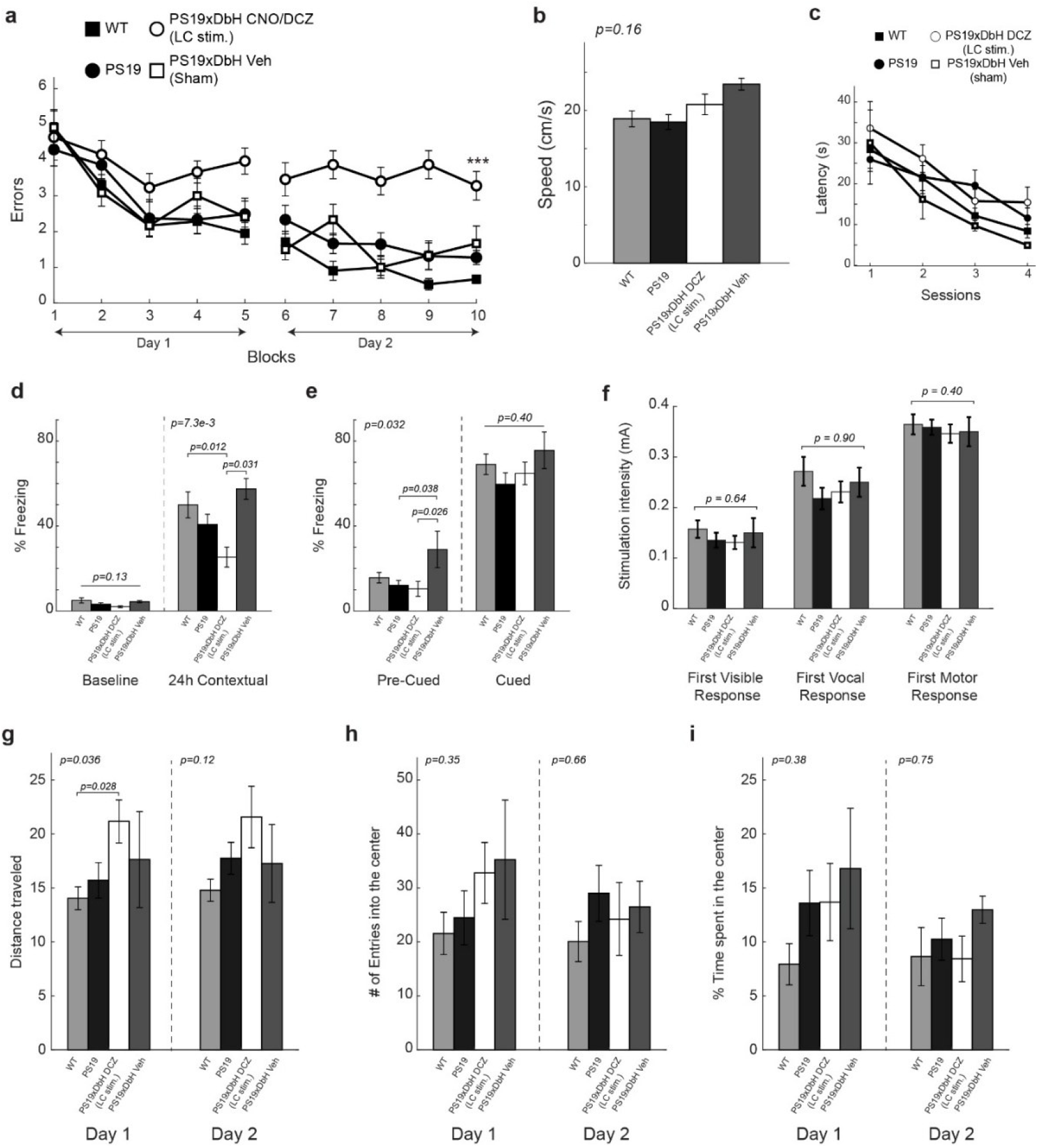
Behavioral performance of experimental mice. **a)** Long-term LC stimulation induced impairment of PS19xDbH RAWM performance. ANOVA for repeated measures among all (day 2): F_(3,44)_ = 19.90, p<0.0001. One-way ANOVA for block 10: F_(3,44)_ = 17.92, p < 0.0001; Tukey-Kramer: p < 0.0001 PS19xDbH LC stim. vs. WT or PS19, p = 0.0276 PS19xDbH LC stim vs. PS19xDbH Veh. WT: n = 14, 6 males, 8 females, PS19: n = 17, 8 males, 9 females, PS19xDbH stim: n = 13, 5 males, 8 females, PS19xDbH Veh: n = 4, 2 males, 2 females. **b-c)** Testing with the visible platform task to assess visual-motor-motivational deficits. Animal data in a) did not show any difference in average speed (ANOVA: F_(3,36)_ = 1.85, p = 0.155) and time to find the visible platform (ANOVA for session 1 to session 4: F_(3,36)_ = 0.54, p = 0.656; F_(3,36)_ = 0.66, p = 0.582; F_(3,36)_ = 0.37, p = 0.775; F_(3,36)_ = 1.32, p = 0.284). WT: n = 14, 6 males, 8 females, PS19: n = 12, 7 males, 5 females, PS19xDbH stim: n = 11, 5 males, 6 females, PS19xDbH Veh: n = 3, 1 male, 2 females. **b)** Long-term LC stimulation induced impairment of contextual memory (24h: ANOVA F_(3,43)_ = 4.56, p = 0.0073, Tukey-Kramer: p = 0.012 PS19xDbH LC stim vs. WT, p = 0.031 PS19xDbH LC stim vs. PS19xDbH Veh) No differences were detected during baseline assessment (ANOVA among all: F_(3,43)_ = 1.98, p = 0.13). WT: n = 14, 6 males, 8 females, PS19: n = 17, 9 males, 8 females, PS19xDbH stim: n = 12, 6 males, 6 females, PS19xDbH Veh: n = 4, 2 males, 2 females. **e)** Freezing responses after the auditory tone were the same among the 4 groups shown in d) in the cued conditioning test. (ANOVA among all: F_(3,43)_ = 1, p = 0.4). PS19×DbH Veh exhibited a higher baseline freezing level before tone presentation, which was likely due to the small sample size (n = 4). (ANOVA among all: F_(3,43)_ = 3.21, p = 0.032, Tukey-Kramer: p = 0.038 PS19xDbH Veh vs. PS19, p = 0.026 PS19xDbH Veh vs. PS19xDbH LC stim). **f)** No difference was shown among the groups in d-e) during assessment for sensory threshold. ANOVA among all: for visible response F_(3,44)_ = 0.57, p = 0.637; for vocal response F_(3,44)_ = 0.19, p = 0.904; for motor response F_(3,44)_ = 1, p = 0.404. WT: n = 14, 6 males, 8 females, PS19: n = 17, 8 males, 9 females, PS19xDbH stim: n = 13, 5 males, 8 females, PS19xDbH Veh: n = 4, 2 males, 2 females. **g-i)** Open field showed PS19xDBH LC stim traveled significantly longer distance at day 1 compared to WT. (F_(3,44)_ = 3.10, p = 0.0364, Tukey-Kramer: p = 0.0284 PS19xDbH LC stim vs. WT). No difference in travel distance was shown at day 2 (F(3,44) = 2.08, p = 0.117). All groups of mice showed a similar number of entries into the center zone (F_(3,44)_ = 1.12, p = 0.35; F_(3,44)_ = 0.54, p = 0.66 for day 1 and day 2, respectively) and similar percentage of time spent in the center zone on both days (F_(3,44)_ = 1.06, p = 0.375; F_(3,44)_ = 0.4, p = 0.754 for day 1 and day 2, respectively). WT: n = 14, 6 males, 8 females, PS19: n = 17, 8 males, 9 females, PS19xDbH stim: n = 13, 5 males, 8 females, PS19xDbH Veh: n = 4, 2 males, 2 females.

We next assessed the effect of chronic LC stimulation on associative memory using the fear conditioning paradigm, which depends on the hippocampus and amygdala (Phillips and LeDoux, 1992) and is commonly impaired in Alzheimer’s disease (Swainson et al., 2001). Baseline freezing levels were comparable across all groups (p=0.13, one-way ANOVA test. **Fig. 2d**). However, PS19 mice receiving LC stimulation showed impaired contextual memory 24 hours post-training, with significantly reduced freezing compared to WT littermates (p=0.012, post -hoc Tukey-Kramer test. p = 7.3e-3, one-way ANOVA test across groups) as well as compared to sham control mice that received vehicle solutions (p = 0.031, post-hoc Tukey-Kramer test. **Fig. 2d**). In the cued fear conditioning test on day 3, which relies on the amygdala but not the hippocampus (Phillips and LeDoux, 1992), although the sham control PS19 mice exhibited slightly higher freezing duration in the pre-cued periods than PS19 mice received chronic LC stimulation and PS19 mice (p=0.026 and p=0.038, respectively, post-hoc Tukey-Kramer tests), there were no significant differences in freezing duration among the four cohorts of mice when the cue tone was played (p = 0.40, one-way ANOVA test. **Fig. 2e**). To ensure that the observed group differences in electric shock–mediated memory tasks were not due to altered nociception, we performed a sensory threshold test prior to the memory tasks. We found no evidence that the treatments affected pain perception, indicating that the differences in the memory tasks likely resulted from chronic LC stimulation. (**Fig. 2f**).

Lastly, we conducted the open field test to assess exploratory and anxiety-like behavior (Acquarone et al., 2019). PS19 mice that received long-term LC stimulation exhibited increased exploratory activity in day 1 compared to WT littermates (p=0. 036 one-way ANOVA test, p=0.028, post-hoc Tukey-Kramer test. **Fig. 2g**). However, the difference vanished in day 2 (p=0.12, one-way ANOVA test. **Fig. 2g**). We failed to observe any significant difference among the four cohorts for both day 1 and day2 in terms of their number of entering the center zone (p=0.35, and p=0.66, respectively, one-way ANOVA test. **Fig. 2h**) or their time spent in the center zone (p=0.38, and p=0.75, respectively, one-way ANOVA test. **Fig. 2i**). Taken together, these results demonstrate that long-term LC stimulation in PS19 mice impairs memory performance but did not increase anxiety-like behavior.

### Chronic LC stimulation impaired the hippocampal LTP in PS19 mice

Given the memory impairments observed only in PS19xDbH mice that received long-term LC stimulation, we next assessed synaptic plasticity by recording long-term potentiation (LTP) in hippocampal slices from PS19xDbH mice that received chronic LC stimulation as well as their WT littermates and PS19 mice.

At both 10- and 30-minute post-tetanus, compared to age-matched WT controls, 7-month-old PS19 mice showed a slight, but non-significant reduction in LTP (p=0.56, and p=0.27, respectively, post hoc Tukey-Kramer test. **Fig. 3**). However, consistent with our RAWM results, PS19xDbH mice that received long-term LC stimulation exhibited the most pronounced impairment, showing the lowest fEPSP slopes among the three groups. One-way ANOVA test confirmed that there is a significant difference in the fEPSP slope among three cohorts of mice (p=0.038, and p=0.029, respectively) at the 10- and 30-minutes post-tetanus but not at the 120-minutes post-tetanus (p=0.07) (**Fig. 3b-d**). Post hoc Tukey-Kramer tests revealed a significant difference between the LC stimulation group and the WT mice group at both 10- and 30-minutes post-tetanus (p=0.032, and p=0.022, respectively. **Fig. 3b&c**). These results indicate that chronic LC stimulation impairs hippocampal LTP in PS19 mice, aligning with the observed behavioral deficits and suggesting a synaptic mechanism underlying the cognitive decline induced by long-term LC activation in a tau mouse model.

**Figure 3.**
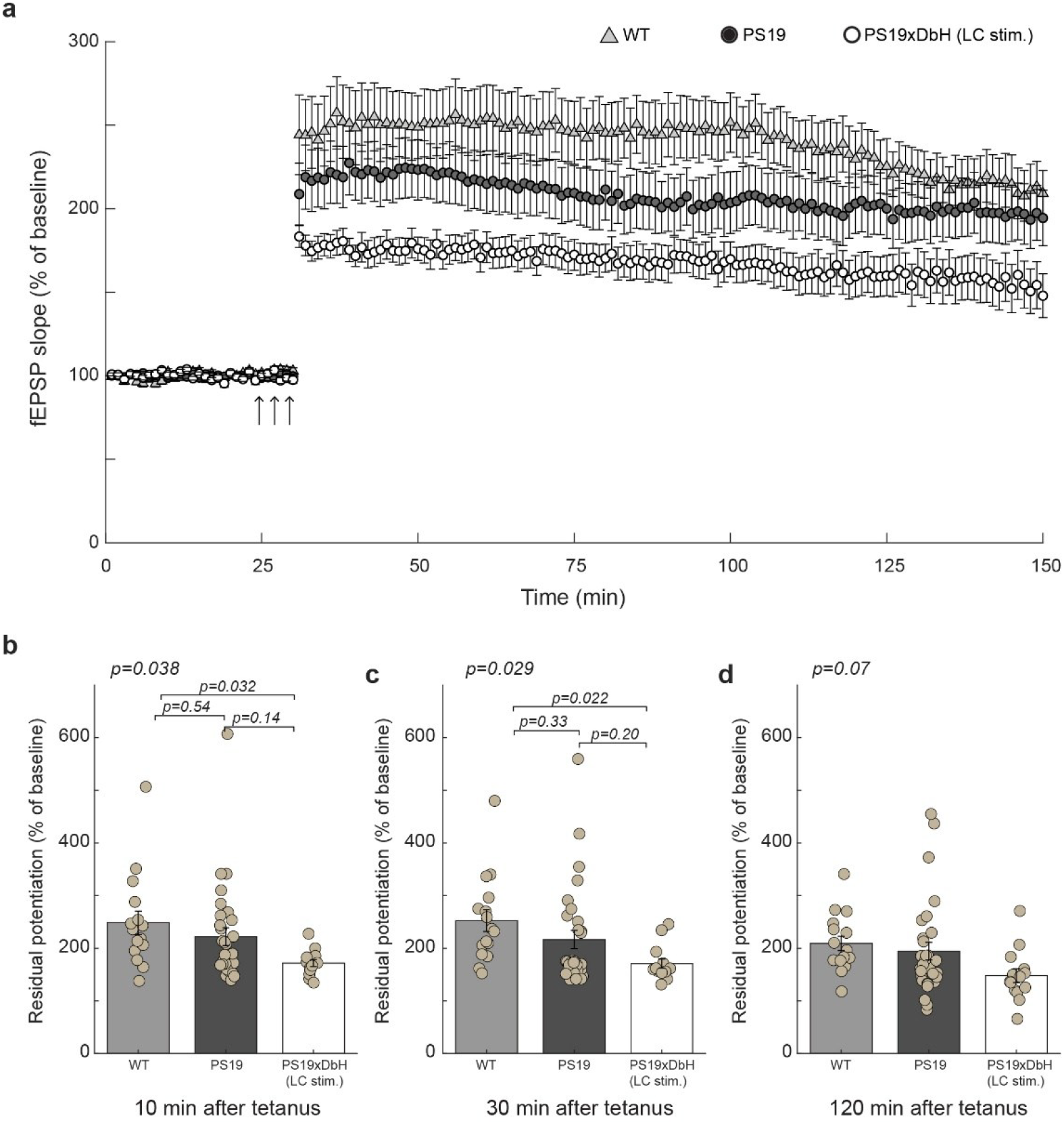
Hippocampal LTP of experimental mice. **a)** Summary graph showing that long-term LC stimulation induced LTP impairment in PS19xDbH LC stim mice. **b-d)** Quantification of the residual potential at 10, 30 and 120 min after tetanus from LTP curves shown in a), respectively. (WT: n = 16 slices/7 animals, 4 males, 3 females; PS19: n = 30 slices/10 animals, 3 males, 7 females; PS19xDbH (LC stim): n = 14 slices/6 animals, 3 males, 3 females). At 10 min, one-way ANOVA among all: F(2, 57) = 3.46, p = 0.038; Tukey-Kramer: p = 0.032 PS19xDbH (LC stim) vs. WT. At 30 min, one-way ANOVA among all: F(2, 57) = 3.75, p = 0.029; Tukey-Kramer: p = 0.02 PS19xDbH (LC stim) vs. WT. At 120 min, one-way ANOVA among all: F(2, 57) = 2.78, p = 0.070.

### Neurodegeneration and neuroinflammation increased in the hippocampus of PS19 mice undergoing chronic LC stimulation

Neuroinflammation is a key contributor to the pathogenesis of AD (Heppner et al., 2015; Kinney et al., 2018). Activated astrocytes and microglia can accelerate disease progression and promote neuronal loss and cognitive decline (Deng et al., 2024; Han et al., 2025). To assess whether chronic LC stimulation increased inflammation levels in the hippocampus, we performed immunostaining for glial fibrillary acidic protein (GFAP), a marker of activated astrocytes (**Fig. 4a**). PS19xDbH mice with chronic LC stimulation exhibited a significantly higher percentage of GFAP-positive area compared to PS19 controls (p = 0.027, Student’s t-test. **Fig. 4b**), indicating elevated astrocytic activation and increased neuroinflammation in the hippocampus. Since elevated neuroinflammation usually results in neurodegeneration, we next examined whether chronic LC stimulation induced neuronal loss in the hippocampus. To this end, we used NeuN staining, a neuronal marker, to evaluate number of neurons in the same slices. As expected, PS19 mice that received chronic LC stimulation showed a significant reduction in NeuN-positive cells compared to PS19 mice (p = 0.03, Mann-Whitney U test. **Fig. 4c**), suggesting a potential increase in hippocampal neuronal loss following long-term LC stimulation.

**Figure 4.**
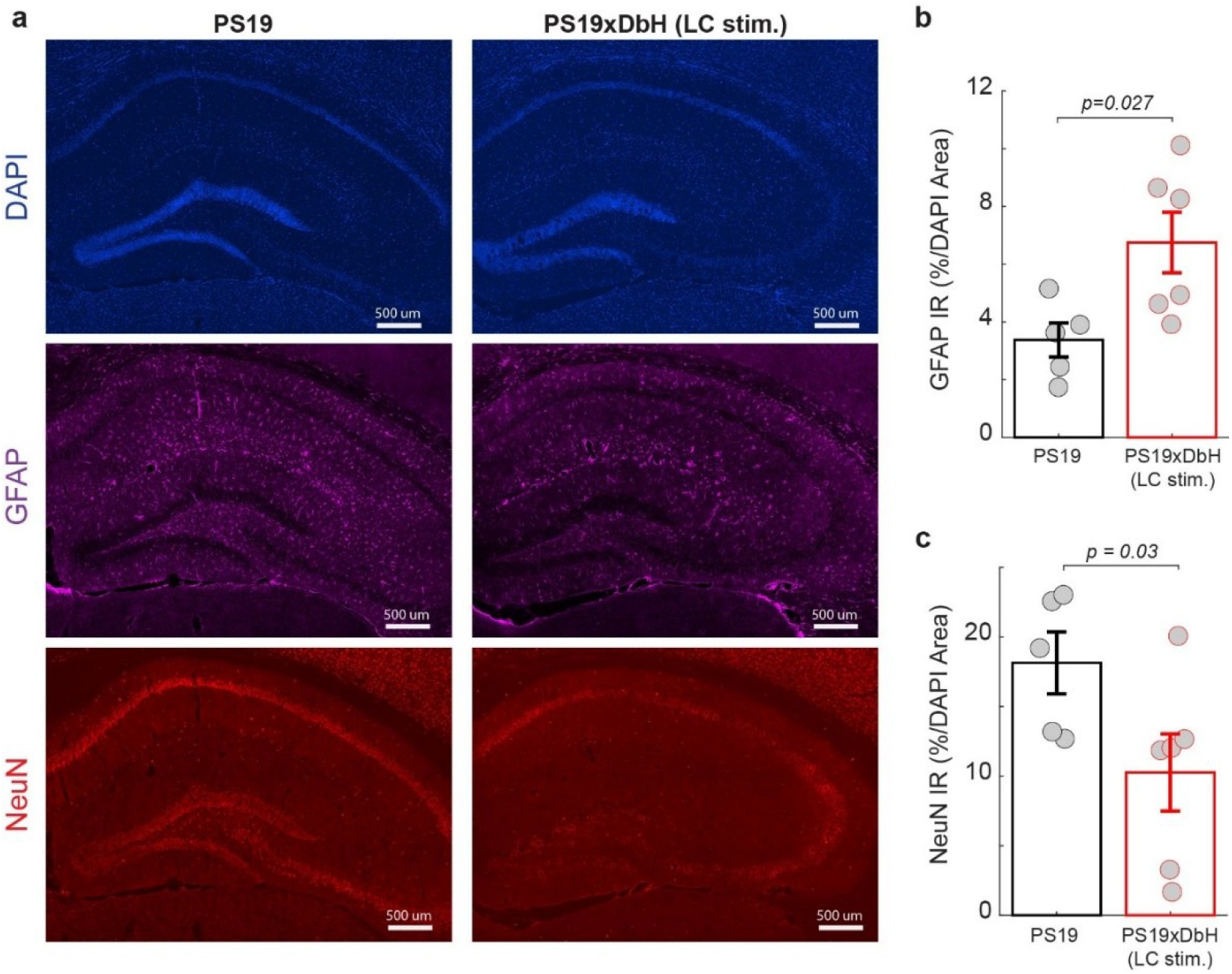
Chronic stimulation increased neurodegeneration and neuroinflammation in the hippocampus. **a)** Example IHC images of DAPI, GFAP and NeuN signals in the hippocampus of PS19 mice with and without LC stimulation. **b)** Percent of overlap of GFAP and DAPI signals in the hippocampus of PS19 mice with and without LC stimulation. **c)** Percent of overlap of NeuN and DAPI signals in the hippocampus of PS19 mice with and without LC stimulation.

### Chronic LC stimulation increased Tau pathology in the LC

To assess Tau pathology in the LC, we examined phosphorylated Tau levels using AT8 immunostaining. AT8 recognizes Tau phosphorylated at Ser202/Thr205, a well-established marker of pathological Tau (Biernat et al., 1992; Goedert et al., 1995). Coronal brainstem sections containing the LC were imaged using confocal microscopy, and LC neurons were identified by their reactivity to DbH antibodies (**Fig. 5a**). We then quantified the percentage of AT8 signal colocalized with DbH-positive neurons in a representative coronal slice for each animal. The representative slice was determined during confocal pre-scanning as the slice that contained the largest number of LC neurons. LC-stimulated PS19xDbH mice exhibited significantly higher levels of phosphorylated Tau in the LC compared to PS19 controls (p = 0.048, Student’s t-test. **Fig. 5b**). Representative images revealed stronger and more widespread AT8 immunoreactivity in the LC of stimulated mice. These results indicate that chronic LC stimulation aggravates Tau pathology within LC neurons in PS19 mice.

**Figure 5.**
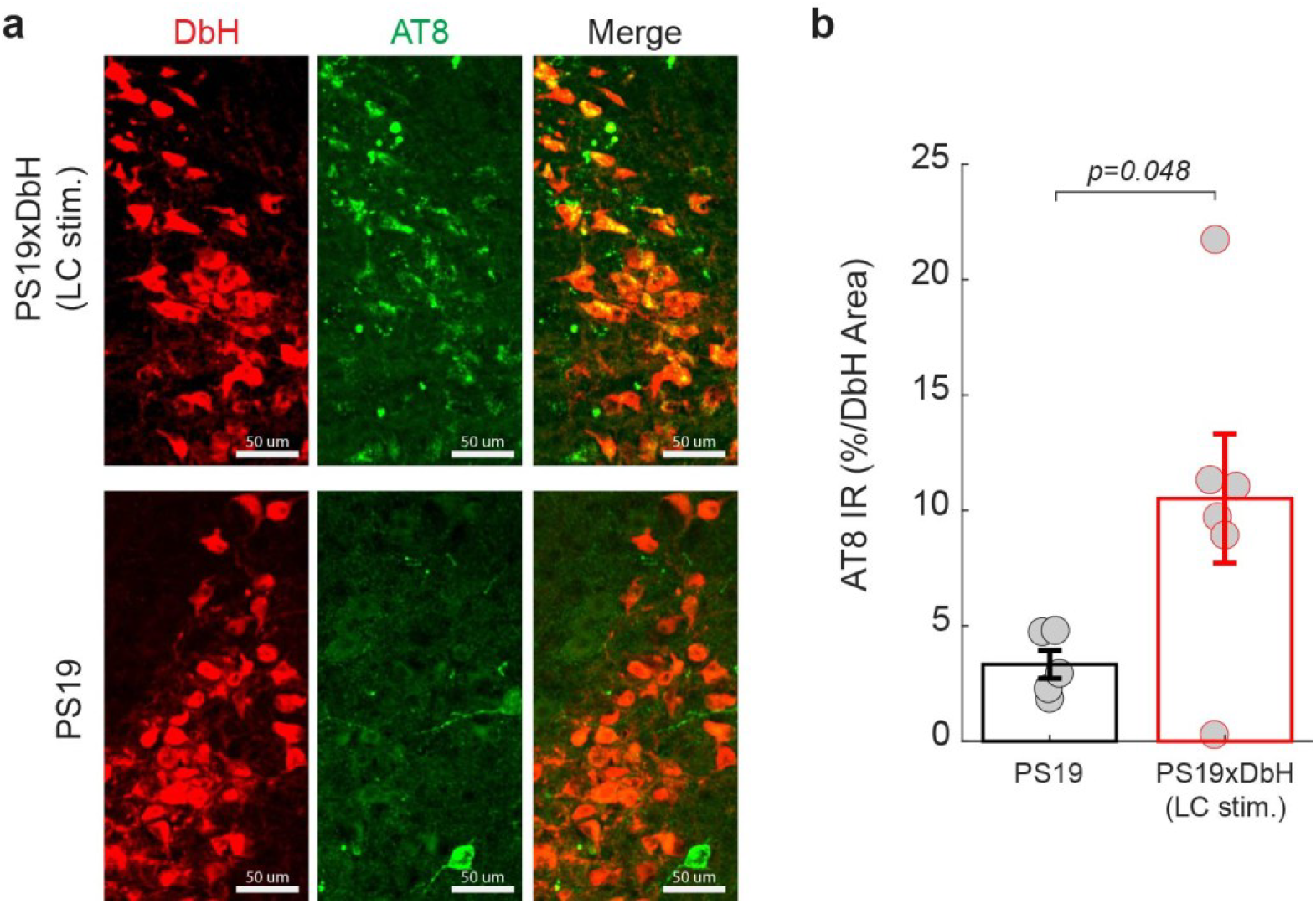
Chronic stimulation increased Tau pathology in the LC. **a)** Example IHC images of TH and phosphorylated Tau (AT8) signals in the LC of PS19 mice with and without LC stimulation. **b)** Percent of overlap of TH and AT8 signals in the LC of PS19 mice with and without LC stimulation.

### Chronic LC stimulation increased AEP expression in the LC

Asparagine endopeptidase (AEP) has been shown to promote Tau pathology by cleaving Tau at residue N368 and facilitating its hyperphosphorylation. Prior studies have reported that the NE metabolite DOPEGAL can activate AEP, contributing to Tau hyperphosphorylation and memory impairment (Kang et al., 2022, 2020). Therefore, the elevated tauopathy may result from increase in AEP expressions. To investigate whether chronic LC stimulation affects AEP activity, we quantified the colocalization of AEP with tyrosine hydroxylase (TH), a marker of LC neurons (**Fig. 6a**). The percentage of AEP-positive signal overlapping with TH-positive cells was significantly higher in LC-stimulated PS19xDbH mice compared to PS19 controls (p = 0.03, Mann-Whitney U test. **Fig. 6b**). Together, these results suggest that long-term LC stimulation upregulates AEP expression in LC neurons, potentially contributing to the exacerbation of Tau pathology.

**Figure 6.**
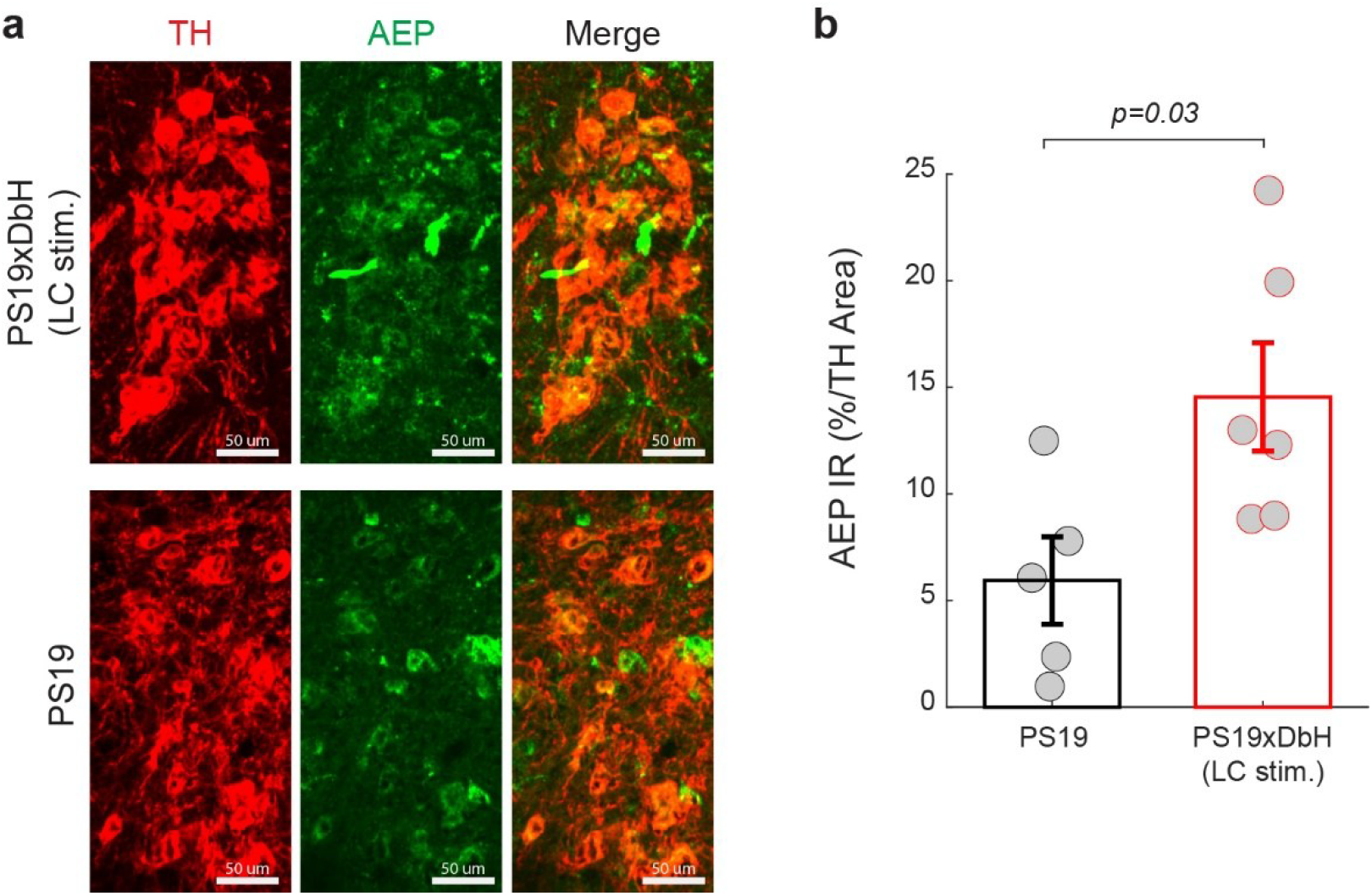
Chronic stimulation increased AEP expression in the LC. **a)** Example IHC images of TH and AEP signals in the LC of PS19 mice with and without LC stimulation. **b)** Percent of overlap of AEP and TH signals in the LC of PS19 mice with and without LC stimulation.

## Discussion

In this study, we investigated the long-term effects of chemogenetic activation of the LC on cognition and disease pathology in PS19 mice, a tauopathy model of AD. Our findings showed that long-term LC stimulation over a three-month period leads to worsening of spatial and contextual memory performance, impaired hippocampal synaptic plasticity, and exacerbated Tau pathology and neuroinflammation. These results suggest that overactivation of the LC-NE system may accelerate tau-related pathology in AD.

Previous studies have shown that short-term activation of the LC can enhance memory performance in a reversal learning task for a TgF344-AD rat model, which expresses both mutant amyloid precursor protein (APPsw) and presenilin-1 (PS1ΔE9) (Rorabaugh et al., 2017). Although this rat model eventually develops Tau pathology at older ages, it is primarily an amyloid-driven model. In this context, transient increases in NE via CNO-mediated activation of DREADD receptors expressed in LC neurons before each behavioral session may have exerted protective effects against Aβ-related toxicity, likely by inhibiting Aβ aggregation and enhancing TrkB signaling (Liu et al., 2015; Zou et al., 2019). On the other hand, long-term elevation of NE has been associated with detrimental effects in the context of Tauopathy. For example, chronic treatment with the NE reuptake inhibitor reboxetine led to worsened memory deficits and neurodegeneration in tauopathy mouse models (Jeong et al., 2025). These findings align with our results, in which chronic LC stimulation in PS19 mice led to a significant decline in spatial working memory and impaired contextual fear memory. Together, these data suggest that the effects of LC activation are both time-dependent and pathology-dependent, while short-term NE elevation may be beneficial in the context of amyloid pathology, long-term activation appears to be harmful in the setting of tau pathology.

The cognitive deficits observed in PS19 mice subjected to long-term LC activation were further supported by electrophysiological evidence of impaired synaptic plasticity. LTP in the hippocampus, a well-established cellular correlate of learning and memory, was significantly reduced in PS19 mice that received long-term LC stimulation as compared to their WT littermates (**Figure 3b&c**). Chronic LC stimulation may have an even more deleterious impact in PS19 mice because they express the human MAPT P301S tau mutation, which causes familial frontotemporal dementia (Ghetti et al., 2015). These findings suggest that chronic LC stimulation in the tau mouse model not only affects behavioral outcomes but also disrupts synaptic function. To investigate potential mechanisms underlying these functional impairments, we examined Tau phosphorylation in the LC using AT8 immunostaining. We observed a significant increase in pTau levels in the LC neurons of stimulated PS19xDbH mice compared to PS19 controls. Since the LC is the earliest region to exhibit pTau accumulation in both humans and rodent models of AD (Braak et al., 2011; Rorabaugh et al., 2017), this result supports the notion that LC overactivity may initiate or exacerbate local Tau pathology. Our data provide in vivo evidence that prolonged LC stimulation alone is sufficient to elevate pTau in this region, without requiring additional factors such as Aβ accumulation (Stancu et al., 2014; Li et al., 2016; Vergara et al., 2019; Zhang et al., 2020).

To further investigate the mechanism underlying Tau pathology, we examined the expression of AEP, an enzyme that cleaves Tau at residue N255/N368 and facilitates its aggregation and phosphorylation (Zhang et al., 2014). AEP can be activated by DOPEGAL, a reactive metabolite produced during NE degradation via monoamine oxidase (Kang et al., 2020, 2022). In our study, AEP expression was significantly elevated in LC neurons following chronic stimulation, suggesting that prolonged NE release may lead to increased DOPEGAL production and subsequent AEP activation. Our results, therefore, offer new evidence for this pathway, which in turn provides a plausible mechanistic link between sustained LC activity, altered NE metabolism, and enhanced Tau pathology, ultimately contributing to synaptic dysfunction and cognitive decline (Kang et al., 2020).

Beyond the LC, we also examined the hippocampus for signs of neuroinflammation and neurodegeneration. We found that astrocyte activation, assessed via GFAP staining, was significantly increased in the hippocampus of LC-stimulated PS19 mice, indicating a heightened inflammatory response. Although NE is generally anti-inflammatory (O’Sullivan et al., 2009; Heneka et al., 2010; O’Donnell et al., 2012), previous studies showed that chronic stress or NE elevation can trigger glial activation and cytokine release (Sugama and Kakinuma, 2021), which in turn contribute to synaptic dysfunction and neuronal vulnerability (Woodburn et al., 2021; Sanguino-Gómez et al., 2022). In addition, NeuN staining in this study revealed a significant reduction in neuronal density in the hippocampus, indicating that prolonged LC activation not only accelerates glial activation but also drives overt neuronal loss. Although future work is necessary to understand the extent to which the observed apoptosis of hippocampal neurons is due to aggregated tau forms or elevated cleaved caspase-3 activity, our findings are consistent with previous reports linking sustained NE elevation to neuronal vulnerability in AD models (Jeong et al., 2025).

Taken together, our findings reveal a complex role for the LC-NE system in AD. While the LC is essential for regulating various perceptual and cognitive functions, excessive or prolonged LC activation may have harmful effects in the context of tauopathy. These results highlight the importance of considering both the timing and duration of neuromodulatory interventions. Although targeting the LC remains an appealing strategy for AD treatment, our data suggest that overactivation of the LC during early disease stages may aggravate rather than improve pathology.

Several limitations of this study should be acknowledged. First, our study focused on one tauopathy mouse model (PS19) and may not generalize to other mouse models used in AD studies, particularly those with Aβ pathology. Second, we used a single dose and duration for chemogenetic LC stimulation. Since it is possible that excessive or prolonged stimulation of the LC contributed to the detrimental outcomes observed here, future work should explore dose-dependent effects and different stimulation durations to determine whether an optimal therapeutic dose exists. Third, while our results point to AEP and pTau accumulation in LC neurons as potential factors, additional studies are needed to directly measure NE and DOPEGAL levels and to confirm their causality in this pathway.

In conclusion, our study provides new evidence that LC overactivation worsens memory deficits, impairs hippocampal LTP, and promotes Tau pathology in a Tau-overexpressing AD model. These findings suggest that while the LC plays a central role in numerous brain functions, the consequences of its activation depend on both disease context and stimulation dose. Prolonged NE elevation in a tauopathy model may worsen pathology, whereas their effect in amyloid-related models may yield different outcomes (unpublished data). Future studies will be important to clarify how LC activity varies across disease stages and pathologies, and to develop treatment strategies that fine-tune LC activity to harness the beneficial effects of NE while avoiding downstream toxicity.

## Acknowledgements

This work was supported by NIH R01AG075114, and R01NS119813.

## Disclaimer

Q.W. is the co-founder of Sharper Sense.

## Data Availability

All data will be available upon request

